# A High-Throughput Bone Marrow 3D Co-Culture System to Develop Resistance to B Cell Receptor Signaling Targeted Agents in B Cell Non-Hodgkin Lymphoma

**DOI:** 10.1101/2025.01.14.632958

**Authors:** Alex Zadro, Alberto Arribas, Maria Vittoria Colombo, Eleonora Cannas, Filippo Spriano, Luciano Cascione, Afua Adjeiwaa Mensah, Federico Simonetta, Dalila Petta, Chiara Arrigoni, Christian Candrian, Matteo Moretti, Francesco Bertoni

## Abstract

B cell receptor (BCR) signaling plays a central role in the pathogenesis of B cell lymphomas, making it a crucial therapeutic target. The advent of BCR-targeted inhibitors, particularly those directed at PI3K and BTK, has revolutionized treatment for B cell non-Hodgkin lymphoma (B-NHL). However, therapeutic resistance remains a significant clinical challenge. Increasing evidence suggests that the tumor microenvironment (TME), particularly the bone marrow (BM) microenvironment, is crucial in driving cancer progression and therapeutic resistance. The BM microenvironment provides a specialized niche where lymphoma cells can evade therapy through interactions with stromal cells and extracellular matrix (ECM) components. Bone marrow stromal cells (BMSCs) contribute significantly to this resistance.

In this study, we developed an *in vitro* 3D model to better understand B cell lymphoma biology and drug resistance mechanisms. We co-cultured lymphoma cell lines with primary BMSCs in a 3D fibrin gel matrix using a high-throughput and automated system. Our results revealed that BMSCs modulate lymphoma cell growth and reduce their sensitivity to the PI3K inhibitor copanlisib and the BTK inhibitor ibrutinib. Furthermore, this model allowed us to identify IGFBP-3, Serpin E1, and PTX-3 as potential mediators of therapeutic resistance. These findings underscore the value of using 3D co-culture models in preclinical settings to more accurately study drug resistance, as they more closely simulate the BM microenvironment’s complexity than traditional 2D models, thus improving the predictive value of drug testing in B cell lymphomas.

**Visual Abstract.**
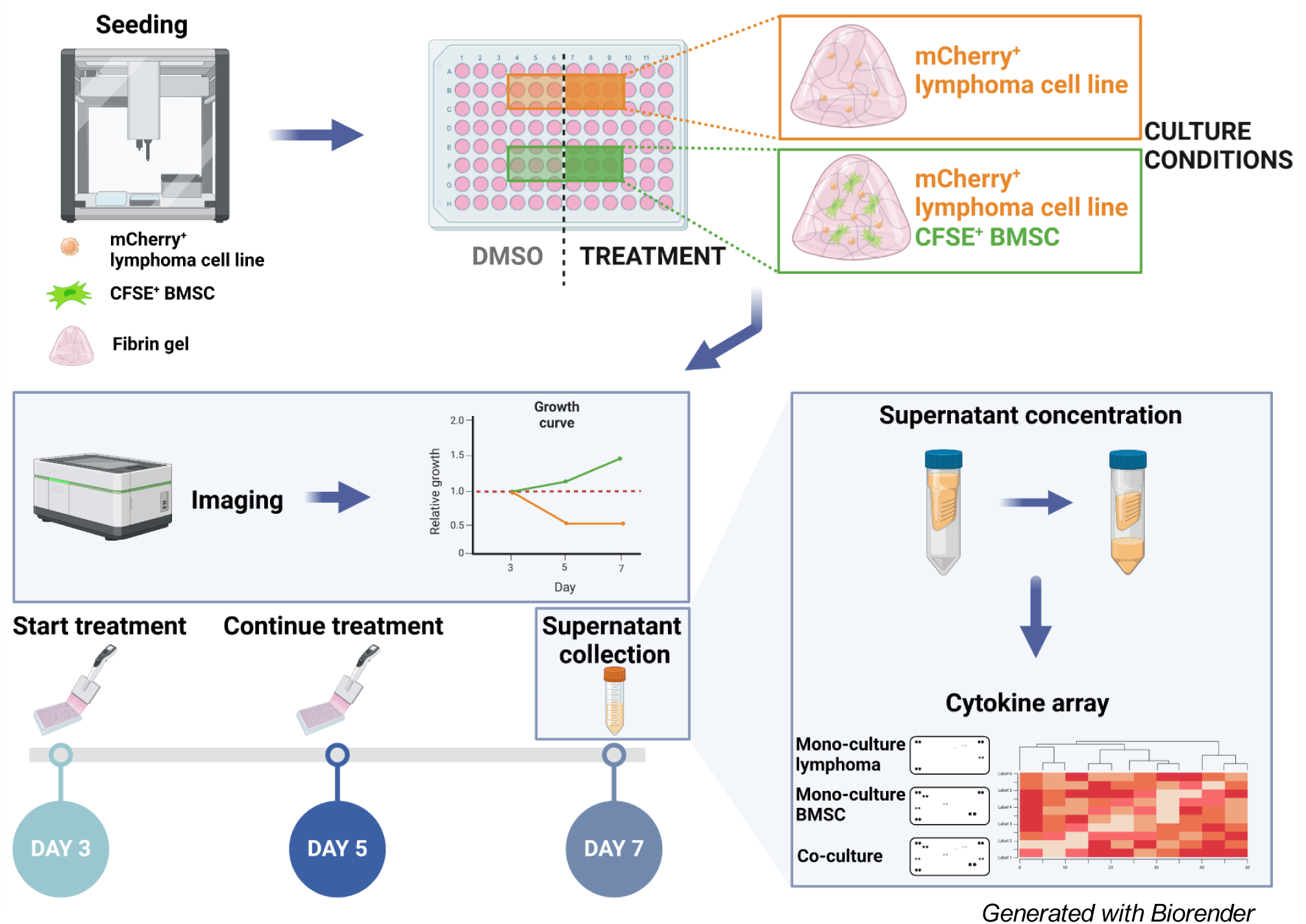

## Introduction

B cell receptor (BCR) signaling is fundamental in driving the pathogenesis of B cell lymphomas, making it a key target for therapeutic intervention^1–4^. The development of BCR-targeted inhibitors, particularly those targeting PI3K and BTK, has significantly expanded treatment options for patients with B cell non-Hodgkin lymphomas (B-NHL)^1–4^. However, the development of resistance to treatments remains a significant clinical challenge in managing B-NHL patients^5–9^.

The tumor microenvironment (TME) is a crucial factor not only in driving cancer progression but also in promoting resistance to therapies. This has been demonstrated in various types of cancer^10,11^ and also in B cell lymphomas^12,13^. Within this context, the bone marrow (BM) microenvironment is critical, acting as a “privileged” niche where cancer cells can evade therapeutic interventions^14–21^.

The BM microenvironment comprises cellular components and a 3D extracellular matrix (ECM) with unique characteristics. The stromal compartment contributes to the pathobiology of multiple hematological malignancies^22^. In particular, bone marrow stromal cells (BMSCs) are multipotent stem cells^23^ that have been implicated in promoting therapeutic resistance in several hematological malignancies by the release of inflammatory cytokines^24^.

In addition to the cellular composition, the ECM within the bone marrow also plays an active role in influencing tumor behavior, histopathology, and therapeutic response^25^. The cancer-associated ECM is not only a structural feature of tumors but also an active driver of cancer progression, which underscores the importance of including a 3D environment in preclinical models to improve the reliability of drug resistance testing in solid tissue environments^26^.

In this study, we established a simplified *in vitro* 3D model to mimic the BM microenvironment in B-NHL, providing a more physiologically relevant platform for investigating lymphoma biology and therapeutic resistance compared to conventional 2D cultures. We co-cultured B cell lymphoma cell lines with primary BMSCs in a 3D fibrin gel matrix with a high-throughput, automated, reproducible seeding, followed by high-content confocal live-cell imaging, enabling real-time monitoring of cellular interactions and drug responses. Our results demonstrate that BMSCs modulate the proliferation and growth dynamics of lymphoma cells and significantly reduce lymphoma cells’ sensitivity to BTK and PI3K inhibitors. Additionally, our model allowed the identification of IGFBP-3, Serpin E1, and PTX-3 as potential novel mediators of therapeutic resistance in B cell lymphomas.

## Material and methods

### Primary cells

The human-derived specimen sampling was conducted under the guidelines, regulations, and procedures of the Ente Ospedaliero Cantonale (Bellinzona, Switzerland) and approved by the local IRB (Approval n. 2020-00029 from Comitato Etico Cantonale). The biological samples used in the study are represented by waste surgical pieces, harvested from patients who signed an informed consent. All the individuals (four females, two males) were subjected to knee replacement surgery and presented OA Kellgren Lawrence grade 3 to 4, evaluated from radiological images. The age of the patients ranged between 58 and 73 years.

### Isolation of primary Bone Marrow Stromal Cells (BMSCs) and culture conditions

Primary BMSCs were isolated from knee biopsies, as previously reported^27^. Following isolation, BMSCs were cryopreserved at −150°C, and all experiments were conducted within a maximum of three weeks after thawing. To create pools of BMSCs, equal numbers of cells from each donor were mixed. The pooled cells were then plated and cultured until they reached passage 3, at which point they were used for experiments.

BMSCs were cultured in MEM Alpha medium (12571063, Gibco) supplemented with 10% fetal bovine serum (FBS; F9665, Sigma-Aldrich), 1% HEPES (15630056, Gibco), 1% Penicillin-Streptomycin (P/S, 10,000 U/mL; 15140122, Gibco) and 5ng/mL basic fibroblasts growth factor (100-18B, Peprotech).

### Cell lines

Details on cell lines are provided in the *Supplementary Appendix*.

### Seeding in 3D fibrin gel

Cell-laden fibrin gels were prepared by suspending lymphoma cells and BMSCs in 4 U/mL thrombin (obtained from the TISSEEL Fibrin Sealant Kit, 025243179, Baxter) diluted in RPMI1640 supplement with L-glutamine, 10% FBS, and 1% P/S. This solution was then mixed with an equal volume of 40 mg/mL fibrinogen (F3879, Sigma) to achieve a final fibrin concentration of 20 mg/mL. After combining the fibrinogen and thrombin, the gels were allowed to polymerize at 37°C for 20 minutes. To maintain sterile conditions, automated high-throughput seeding was performed using the OT-2 liquid handler (Opentrons) equipped with a HEPA module. Cells were seeded into a final volume of 50 μL fibrin gels in Nunc MicroWell 96-Well plates (167008, Thermo Scientific).

### Treatment and cell viability evaluation in 3D fibrin gel

The mCherry-VL51 cells were seeded at 0.075 x 10^6^ cells/mL final concentration, the mCherry-K1718 cells were seeded at 0.5 x 10^6^ cells/mL, the mCherry-SSK41 cells were seeded at 0.25 x 10^6^ cells/mL, the REC-1 and TMD8 cells were seeded at 0.15 x 10^6^ cells/mL. All models were cultured either alone or in co-culture with primary BMSCs, which had been pre-labeled using CFSE (CFSE Cell Labeling Kit, ab113853, Abcam), at 0.75 x 10^6^ cells/mL. Cells were maintained in RPMI-1640 medium supplemented with L-glutamine, 10% FBS, and 1% P/S, with a medium exchange performed on day 1. At day 3, specific treatment conditions, as indicated in the respective figures, were initiated. Copanlisib and ibrutinib were purchased from Selleckchem (TX, USA) and prepared as stock solutions (10mM for copanlisib, 50mM for ibrutinib) in DMSO. On day 5, fresh treatment conditioned medium was added. Live imaging of the cultures was performed on days 3, 5, and 7 using the Opera Phenix Plus High-Content Screening System (PerkinElmer) while keeping the plate at 37°C, 5% CO_2,_ and optimal humidity. The treatment effects were assessed through both morphological observations and quantitative image analysis. The growth curve for each condition was generated by measuring the total mCherry+ area from the maximum projection images, normalizing it first to the respective well at day 3, and then to the DMSO control for each condition. Additionally, at day 7, nuclei were counterstained using Hoechst 33342 (B2261, Sigma), and treatment effects were evaluated based on cell counts. Cluster size was determined by quantifying the mCherry+ area, with the data representing the average of the fourth quartile from the cluster size distribution for each condition.

### Cytokine array

Supernatants were collected from the different 3D experimental conditions at day 7 or after three days of culture (baseline expression) and were concentrated using the Amicon Ultra centrifugal filter units (Z648027, Merck). A 10X concentration was achieved by centrifugation at 16,000 rpm for 1h at 4°C. The concentrated media were then analyzed for 105 soluble human proteins using the Human XL Cytokine Array Kit (ARY022B, Biotechne) according to the manufacturer’s protocol. The signals from the array membranes were captured using the Fusion Solo S imaging system (Vilber) and subsequently quantified using the Protein Array Analyzer plugin in ImageJ (NIH)^28^. The resulting data were normalized to the positive control signals on each membrane, and heatmaps were generated to represent cytokines’ relative secretion. Heatmap visualization was performed using the ComplexHeatmap package in R^29^, with values representing the averaged normalized signal for each cytokine across the experimental conditions.

### MTT assay

Cells were incubated overnight (ON) in 1% FBS medium. Following incubation, cells were seeded into 96-well plates at the densities indicated for each cell line in *Supplementary Table 1*. Cells were cultured in either specific cytokine- or PBS-conditioned in 10% FBS media and stimulated at designated time points, with cytokine concentrations and stimulation durations detailed in *Supplementary Table 1*. All recombinant cytokines used in the experiments were purchased from ProSpec-Tany Technogene (Israel). After stimulation, plates were centrifuged at 1,200 rpm for 5 minutes to remove the conditioned media, and the cell pellets were resuspended in either DMSO or drug-conditioned, phenol red-free media. The specific drug and its concentration for each cell line are indicated in the figures and *Supplementary Table 2*. Cells were incubated under these conditions for 72 hours followed by an MTT assay as previously described^30^.

### Incucyte experiments

Cells were incubated ON in 1% FBS medium. On day 2, cells were transferred to 10% FBS media conditioned with a combination of cytokines, consisting of 100 ng/mL each of IGFBP-3, Serpin E1, and PTX-3, or to PBS-conditioned medium as a control, and incubated for 24 hours. On day 3, cells were resuspended in either DMSO (control) or ibrutinib-conditioned media at the concentrations indicated in the *Supplementary Table 2*. Cells were then seeded into Incucyte-compatible 96 well plates. The plates were imaged every 8 hours for a total of 96 hours using the Incucyte live-cell imaging system (Sartorius) to monitor cell proliferation and growth. The growth curve data were based on the cell area confluence measurements at each time point. Data were normalized to the 8-hour time point.

### Cytokines receptors immunofluorescence

The IbiTreat µ-Slide 8 Well (80826, Ibidi) and IbiTreat µ-Slide 18 Well (81816, Ibidi) were pre-coated by applying poly-L-ornithine solution (A-004-C, Sigma) to each well and incubating at room temperature (RT) for 1 hour. After coating, the solution was removed, and the plates were allowed to air dry. Cells were then seeded in 1% FBS medium at 1 x 10^5^ cells per well (100 µL of a 1 x 10^6^ cells/mL suspension) and allowed to attach ON in the incubator. The media was removed the following day, and the wells were gently washed once with DPBS with calcium and magnesium (14040117, Gibco). Cells were fixed with 4% paraformaldehyde (PFA, 1004965000, Merck) for 15 minutes at RT, followed by three washes with DPBS, each lasting 5 minutes. To block non-specific binding, 4% bovine serum albumin (BSA) in DPBS was added and incubated for 1 hour at RT while shaking. After blocking, the BSA solution was removed, and primary antibodies (more details in *Supplementary Table 3*), diluted in 4% BSA, were added for overnight incubation at 4°C. The next day, wells were washed three times with DPBS for 5 minutes. Secondary antibodies (more details in *Supplementary Table 3*), diluted in 4% BSA with 1:1000 NucBlue Fixed Cell ReadyProbes Reagent (R37606, Invitrogen), were added and incubated for 1 hour at RT. Following secondary antibody incubation, wells were washed three times with DPBS for 5 minutes. Samples were then stored in DPBS containing 0.1% sodium azide (71289-50G, Sigma Aldrich) for long-term preservation at 4°C.

### Graphical representation and statistical analysis

Graphical representation and statistical analyses were performed using GraphPad Prism version 8.01 (https://www.graphpad.com), unless stated otherwise. The specific statistical tests used are detailed in the respective figure legends.

## Results

### 3D co-culture with BMSCs influences the growth pattern of B cell lymphoma cells and reduces their sensitivity to pharmacological inhibition of PI3K

The BM is a privileged niche where lymphoma cells can evade chemotherapy^31^. To gain a deeper insight into the influence of the BM TME on B-NHL therapeutic resistance to BCR signaling inhibitors, we developed a high-throughput 3D BM system. The marginal zone lymphoma (MZL) cell line VL51 was cultured in 3D fibrin gel alone or with primary BMSCs. The automated seeding procedure was highly reproducible, as shown by the uniform cell distribution among wells (Supplementary Figure 1A) and similar measured area values distributions between the VL51 mono- and co-culture conditions (Supplementary Figure 1B). Particularly, the automated seeding procedure uniformly distributed similar numbers of cells among wells with a coefficient of variation of less than 10%, and that was not affected by culture conditions (Supplementary Figure 1C). Treatment with the PI3K inhibitor copanlisib, which has shown promising clinical activity in MZL patients^32^, was initiated on day 3. The growth pattern of VL51 differed based on the various culture conditions. Co-culture with BMSCs induced proliferation, increased the aggregation of VL51 cells at days 3, 5, and 7 (Figure 1A, day 3 images in Supplementary Figure 1D), and significantly decreased the sensitivity to copanlisib (p=0.025, Figure 1B). In the mono-culture, VL51 cells exhibited 20% and 40% reduced proliferation upon copanlisib treatment on days 5 and 7, respectively. In contrast, the viability of VL51 cells was not affected by copanlisib treatment at any analyzed time point in the co-culture. These findings were corroborated by the cell count on day 7 (Supplementary Figure 1E). Furthermore, when left untreated, VL51 cells formed significantly larger clusters (p < 0.001) in the presence of BMSCs (Figure 1C). However, when treated with copanlisib, VL51 cells formed smaller and more dispersed clusters (Figure 1A), comparable in size to those in mono-culture (Figure 1C). These findings indicate that BMSCs influence growth patterns and reduce the copanlisib sensitivity of VL51 cells.

**Figure 1.**
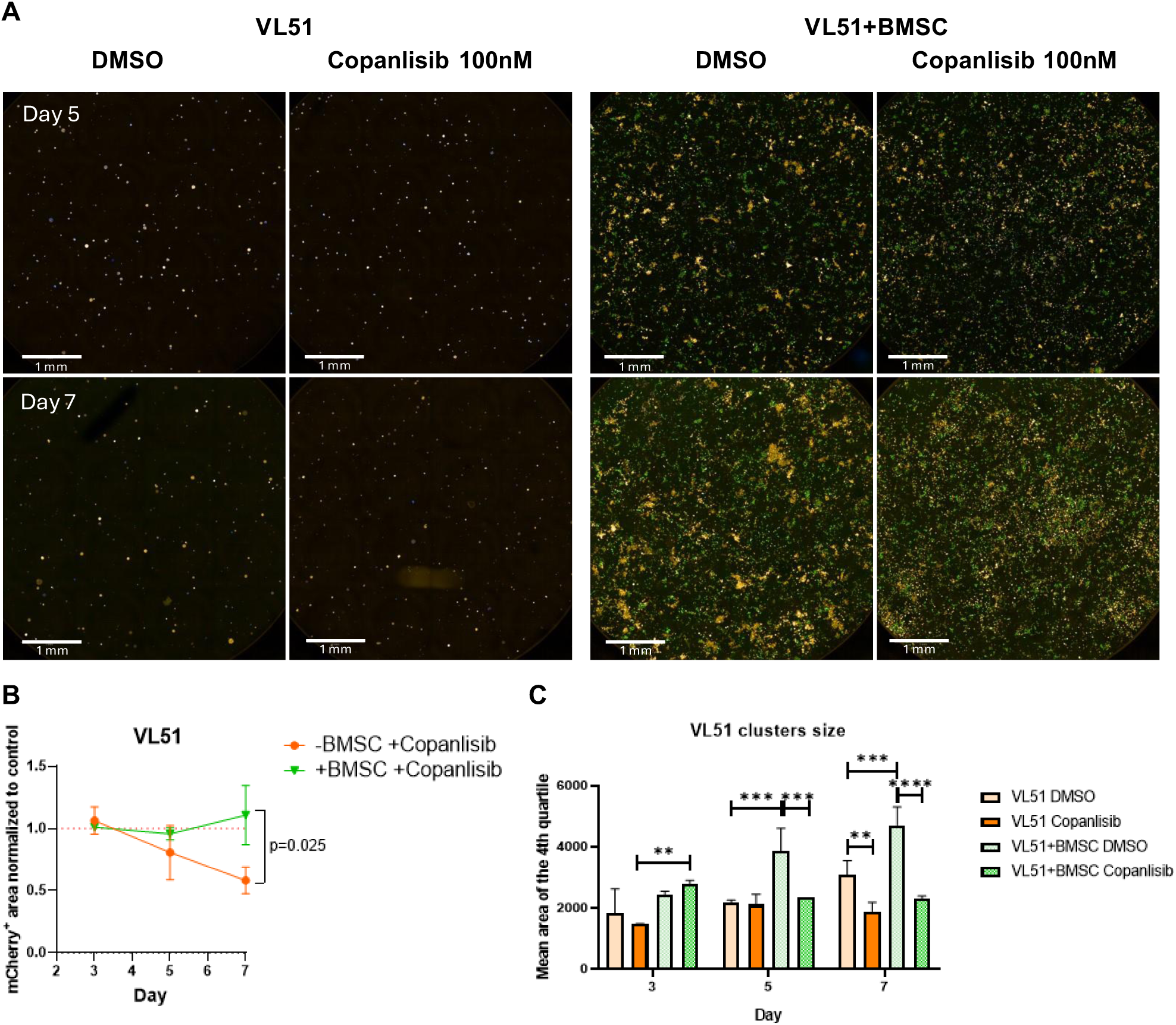
3D co-culture with BMSCs influences the growth pattern of B cell lymphoma cells and reduces their sensitivity to pharmacological inhibition of PI3K. **(A)** Representative immunofluorescence maximum projection images of VL51 (in orange) cultured in the presence or absence of BMSCs (in green) upon DMSO or 100nM copanlisib treatment on day 5 and 7. Images taken with 10x Air objective NA 0.3, WD 5.2mm of the Opera Phenix Plus High-Content Screening System while keeping the plate at 37°C, 5% CO_2,_ and optimal humidity. **(B)** Quantification of the mCherry+ area on maximum projection images, representative of VL51 occupied volume. The measurements shown were normalized to the day 3 time point and the mono- and co-culture treated conditions were normalized to respective DMSO control. Values are plotted as mean with standard deviation of at least three independent biological replicates. Statistical significance tested with Multiple t test. **(C)** Quantification of VL51 cluster size on maximum projection images. The barplot shows the averaged values with standard deviation in the 4^th^ quartile of the distribution of the measured areas of three independent biological replicates. Statistical significance tested with Two-way ANOVA + Dunnett’s multiple comparisons test (**\*** = p < 0.05, **\*\*** = p < 0.01, **\*\*\*** = p < 0.001).

### IGFBP-3, Serpin E1, and PTX-3 reduce the sensitivity of VL51 to copanlisib

We hypothesized that the observed reduced sensitivity of VL51 cells to copanlisib in the presence of BMSCs might be sustained by secreted factors. To identify the potential cytokines involved in this process, we cultured VL51 cells and BMSCs either in mono-culture or co-culture and treated them with DMSO or copanlisib at 100nM. On day 7, we collected the supernatant and analyzed it for the presence of 105 possible secreted factors (*Supplementary Table 4*). We identified 14 cytokines differentially secreted in the different cultures and treatment conditions. Among these, we identified cytokines that were already reported in literature to be involved in therapeutic resistance in lymphoma such as IL-6^33–36^ and VEGF^37,38^ and in homing of malignant cells to the bone marrow such as CXCL12^39^. Moreover, all the identified cytokines were already secreted by BMSCs in the mono-culture, suggesting that BMSCs are the primary source of these cytokines (Figure 2A, whole panel in Supplementary Figure 2A and membranes in Supplementary Figure 2B). Cytokines presenting an increased secretion in the co-culture conditions upon copanlisib and that had not been very well characterized for their biologic role in the lymphoma field were selected for further analyses. IGFBP-3, CHI3L-1, Serpin E1, PTX-3, VCAM1, and DKK-1 were evaluated as recombinant factors to stimulate VL51 cells before treatment with either DMSO or 10nM copanlisib, and cell viability was assessed using the MTT assay. Among the six cytokines, IGFBP-3, Serpin E1, and PTX-3 significantly reduced VL51 sensitivity to copanlisib (Figure 2B), indicating their possible role in reducing sensitivity to copanlisib in the 3D model. Previous studies have identified TMEM219 and TGFβR1 as the receptors for IGFBP-3^40,41^, uPAR (also known as PLAUR) as the receptor for Serpin E1^42^, and TLR4 as a receptor for PTX-3^43^. Immunofluorescence experiments confirmed clear expression of all receptors in VL51 cells (Figure 2C, secondary antibody controls in Supplementary Figure 2C), further sustaining the potential involvement of IGFBP-3, Serpin E1, or PTX-3 in decreasing the sensitivity to copanlisib in MZL cells.

**Figure 2.**
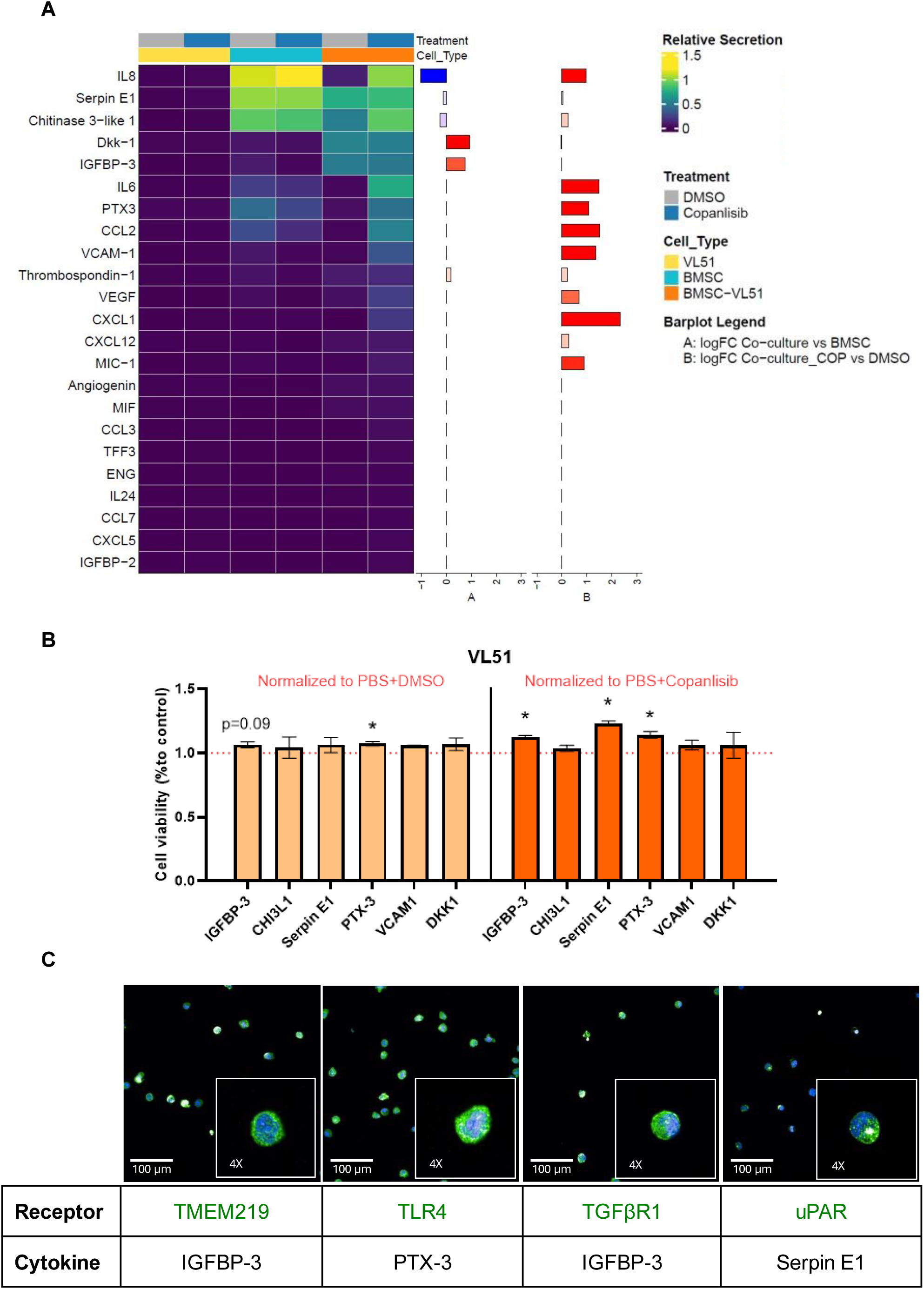
IGFBP-3, Serpin E1, and PTX-3 reduce the sensitivity of VL51 to copanlisib. **(A)** Plot showing a subset of cytokines analyzed with the cytokine array. The heatmap was generated with the relative secretion values (row values normalized to internal positive and negative controls) of each cytokine in the different culture conditions. The bar plots show the log fold change values comparing co-culture to mono-culture upon DMSO (A) or the copanlisib to the DMSO conditions in the co-culture (B). **(B)** Bar plot representing the viability of VL51 cells tested with MTT assay under the different culture conditions. The cytokines-stimulated conditions upon DMSO normalized to the respective PBS+DMSO control, representative of the cytokine-driven proliferative advantage, are plotted in light orange. The cytokines-stimulated conditions upon copanlisib normalized to the respective PBS + copanlisib control, representative of the cytokines’ influence on copanlisib sensitivity, are plotted in dark orange. The plotted values represent the mean with a standard deviation of three independent biological replicates. Statistical significance tested with Multiple t test (**\*** = p < 0.05, **\*\*** = p < 0.01, **\*\*\*** = p < 0.001). **(C)** Representative immunofluorescence images of VL51 cells. In green is shown the expression of the specified receptor, in blue the nuclei. Images taken with 40x Water objective NA 1.1, WD 0.62mm of the Opera Phenix Plus High-Content Screening System at room temperature.

### IGFBP-3, Serpin E1, and PTX-3 reduce the sensitivity to ibrutinib

To extend the potential clinical relevance of our findings, we tested whether IGFBP-3, Serpin E1, and PTX-3 could also reduce sensitivity to BTK inhibitors, another class of BCR signaling inhibitors, largely approved for the treatment of patients with various lymphoma subtypes^2^. Here, we selected *in vitro* models with high sensitivity to BTK inhibitors and derived from lymphomas in which BTK inhibitors are clinically relevant^2^. We studied six models, representative of MZL (Karpas1718 and SSK41), activated B cell-like (ABC) diffuse large B cell lymphoma (DLBCL) (OCI-Ly10 and TMD8), and mantle cell lymphoma (MCL) (REC-1 and MINO). Analysis of the secretome by cytokine array showed that none of the cell lines secreted IGFBP-3, Serpin E1, or PTX-3 at basal conditions (Figure 3A, whole panel in Supplementary Figure 3A, and membranes in Supplementary Figure 3B). Conversely, all the models expressed the corresponding receptors TMEM219, TLR4, TGFβR1, and uPAR (Figure 3B, secondary antibody controls in Supplementary Figure 3C). All cell lines were then stimulated with IGFBP-3, Serpin E1, or PTX-3. Cell viability was measured by MTT assay after 72hrs of exposure to DMSO as a negative control or to equipotent concentrations of the first-generation BTK inhibitor ibrutinib among cells. Stimulation with IGFBP-3, Serpin E1, or PTX-3 did not confer a proliferative advantage to the cells (Figure 4A, light orange bars). However, a general trend of decreased sensitivity to ibrutinib was observed in all lines except for OCI-Ly10, which showed decreased viability. A statistically significant reduction in ibrutinib sensitivity was observed in SSK41 upon IGFBP-3 stimulation, in TMD8 upon IGFBP-3, Serpin E1, and PTX-3, and in MINO when stimulated with Serpin E1 (Figure 4A, dark orange bars).

**Figure 3.**
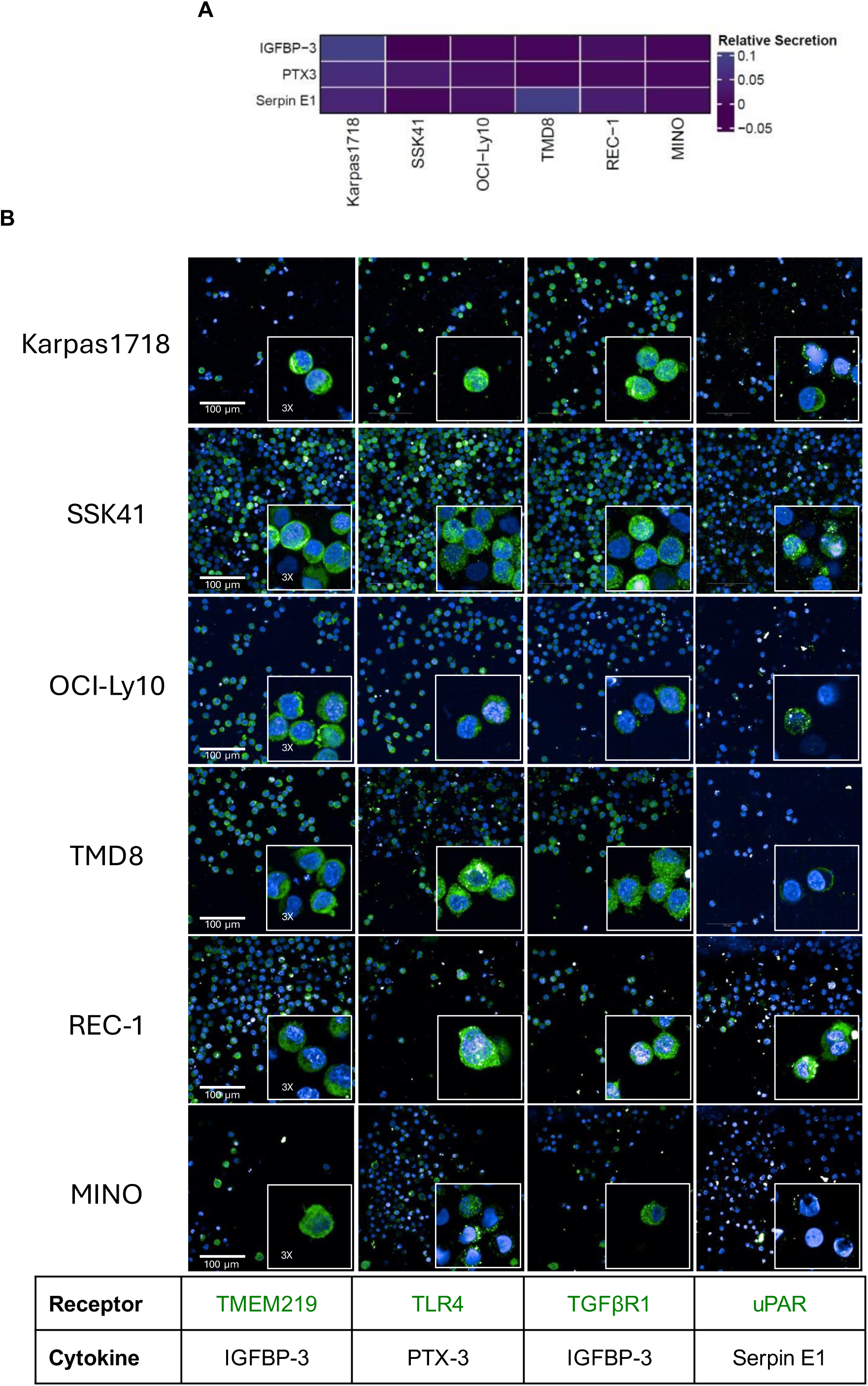
B cell lymphoma cell lines characterization for baseline cytokines and receptors expression. **(A)** Plot showing IGFBP-3, PTX-3, and Serpin E1 secretion analyzed with the cytokine array. The heatmap was generated with the relative secretion values (row values normalized to internal positive and negative controls) of each cytokine in the indicated cell lines. **(B)** Representative immunofluorescence images of the indicated cell lines. In green, it is the expression of the specified receptor, and, in blue, the nuclei. Images taken with 40x Water objective NA 1.1, WD 0.62mm of the Opera Phenix Plus High-Content Screening System at room temperature.

**Figure 4.**
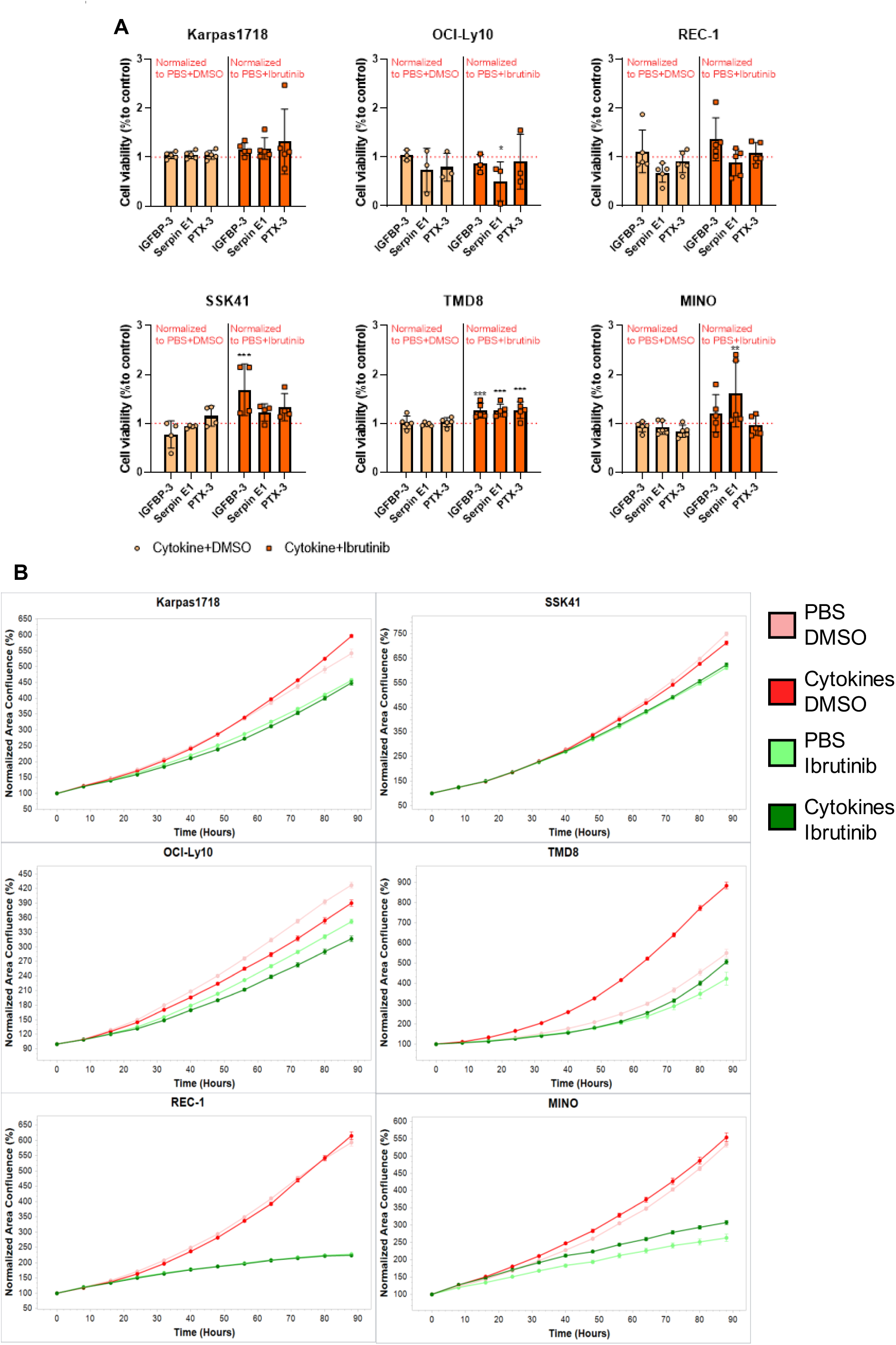
IGFBP-3, Serpin E1, and PTX-3 reduce the sensitivity to ibrutinib. **(A)** Bar plot representing the viability of indicated cell lines tested with MTT assay under the different culture conditions. The cytokines stimulated conditions upon DMSO normalized to the respective PBS+DMSO control, representative of the cytokine-driven proliferative advantage, are plotted in light orange. The cytokines-stimulated conditions upon ibrutinib normalized to the respective PBS+ibrutinib control, representative of the cytokines’ influence on ibrutinib sensitivity, are plotted in dark orange. The plotted values represent the mean with standard deviation of at least three independent biological replicates. Statistical significance tested with Two-way ANOVA + Dunnett’s multiple comparisons test (**\*** = p < 0.05, **\*\*** = p < 0.01, **\*\*\*** = p < 0.001). **(B)** Growth curve of the specified cell lines under the indicated conditions generated with the 20x Air Objective in the Incucyte at 37°C, 5% CO_2,_ and optimal humidity. Plots represent one biological replicate.

We next examined the effects of the combination of IGFBP-3, Serpin E1, and PTX-3, to better mimic the co-culture of lymphoma and BMSCs. Cell lines were concomitantly stimulated with the three cytokines (fixed doses), followed by treatment with DMSO or ibrutinib. Cell proliferation was assessed by live cell imaging, with cells observed for up to 90hrs. Adding all the cytokines provided a proliferative advantage in TMD8 cells while causing a proliferative disadvantage in OCI-Ly10 (Figure 4B). In addition, exposure to the cytokine combo reduced ibrutinib response in MINO and, to a lesser extent, in TMD8. Consistent with the MTT assay results, OCI-Ly10 cells showed a trend towards reduced proliferation and increased sensitivity to ibrutinib upon stimulation with the cytokine combo. The discordant results observed in OCI-Ly10 may result from the limitations of the *in vitro* setting (fixed and single doses, suboptimal exposure time, mono-culture). Altogether, these results suggest that IGFBP-3, Serpin E1, and PTX-3 can confer resistance to BTK inhibitors, although the relevance of these cytokines may vary across cell subtypes or tissues, highlighting the complexity of cytokine-induced resistance mechanisms.

### Anti-tumor activity of BTK inhibitors is reduced when lymphoma cells are grown in 3D co-cultures with BMSCs

Finally, to investigate whether BMSCs contribute to therapeutic resistance against BTK inhibitors, we cultured the lymphoma cell lines VL51, Karpas1718, SSK41, TMD8, and REC-1 in 3D fibrin gel, either in the absence or presence of primary BMSCs. Treatment with the BTK inhibitor ibrutinib (specific concentration indicated in Supplementary Figure 4A-C) was initiated on day 3. On day 7, in the presence of BMSCs, we observed a trend toward increased proliferation in Karpas1718, SSK41, TMD8, and REC-1 (Figure 5A, images in Supplementary Figure 4A-C). Moreover, in the presence of BMSCs, we observed reduced ibrutinib sensitivity in VL51 (p=0.013), and a similar trend, yet not significant, in the other models Karpas1718 (p=0.123), SSK41 (p=0.149), TMD8 (p=0.217) (Figure 5B). Despite using ibrutinib at a concentration considerably higher than the IC_50_ determined in 2D cultures (*Supplementary Table 5*), no significant influence of primary BMSCs on the REC-1 cell line was observed, as ibrutinib had no effect in REC-1 3D mono-culture (Figure 5B). Notably, higher concentrations of ibrutinib were required for all cell lines in the 3D model compared to the 2D culture, suggesting that the presence of a 3D environment and the gel matrix may reduce the sensitivity of cell lines to drug treatment. This observation aligns with previous studies that have reported discrepancies in drug responses between 2D and 3D models^44,45^. Despite these methodological limitations, the findings indicate that BMSCs significantly contribute to mediating resistance to BTK inhibitors in lymphoma.

**Figure 5.**
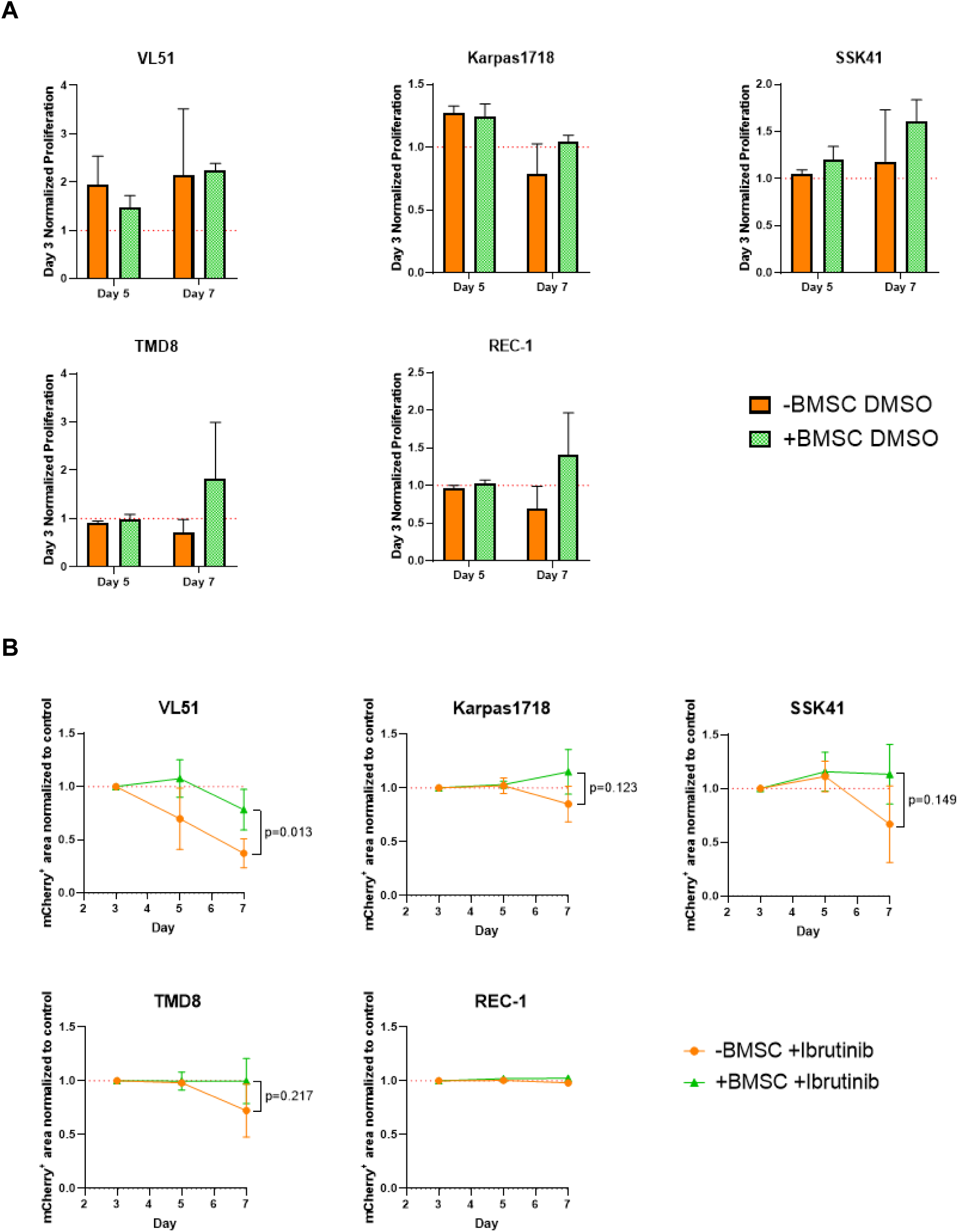
Anti-tumor activity of BTK inhibitors is reduced when lymphoma cells are grown in 3D co-cultures with BMSCs. **(A)** Quantification of the mCherry+ area on maximum projection images, representative of the volume occupied by lymphoma cells. The bar plots show the averaged values with the standard deviation coming from the mono- and co-cultured conditions upon DMSO on day 5 and day 7 normalized to day 3 of at least three independent biological replicates. Statistical significance tested with Multiple t test. **(B)** Quantification of the mCherry+ area on maximum projection images, representative of the volume occupied by lymphoma cells. The measurements shown were normalized to the day 3 time point and the mono- and co-cultured treated conditions were normalized to respective DMSO control. Values are plotted as mean with a standard deviation of at least three independent biological replicates. Statistical significance tested with Multiple t test.

## Discussion

In this study, we developed a simplified *in vitro* 3D BM model for B-cell lymphomas using a high-throughput, automated, and reproducible approach. The BM has long been recognized as a “privileged” niche, providing a protective environment for lymphoma cells to escape chemotherapy^14,16,31^. Our findings extend this concept, demonstrating that our simplified *in vitro* 3D BM model also reduces lymphoma cells’ sensitivity to BCR-targeted agents. In particular, this study provides direct evidence of the role of BMSCs in reducing lymphoma sensitivity to PI3K and BTK inhibitors.

BMSCs emerged as key contributors to the development of therapeutic resistance in our model, consistent with previous studies showing their role in drug resistance in hematological malignancies^21^. Various mechanisms have been reported in the literature to explain this phenomenon. For example, BMSCs can induce resistance through the Notch/Jagged signaling axis in B-cell acute lymphoblastic leukemia (B-ALL)^46^ and activate the NF-kB pathway via VCAM1/VLA4 and VLA5 interactions in ALL^47,48^. BMSCs also release cytokines like CXCL12, which promotes the homing of leukemic cells to the BM by binding to CXCR4, activating pro-survival pathways^39^ such as JAK/STAT in T cells^49^, Wnt/β-catenin in ovarian cancer^50^, and ERK1/2 in hepatocellular carcinoma^51^. Moreover, BMSCs produce IL-6, a cytokine that upregulates anti-apoptotic proteins like MCL-1 and BCL-XL, contributing to drug resistance in multiple myeloma (MM) ^52,53^, as well as in various lymphoma models^33–36^. Consistent with these findings, we confirmed some well-described cytokines involved in lymphoma therapeutic resistance to be secreted in our 3D co-culture model, namely IL-6 and CXCL12. Moreover, using our 3D co-culture system, we identified novel cytokines (IGFBP-3, PTX-3, and Serpin E1) that might promote resistance to lymphoma treatments. Although we did not perform specific killing experiments to confirm that BMSCs exclusively secrete these cytokines, their presence already in BMSC mono-culture, but not in lymphoma mono-cultures, suggests that BMSCs were the primary source. Additionally, we cannot exclude the possibility that BMSCs may influence the sensitivity of lymphoma cell lines to BCR-targeted agents through other secreted factors that were not thoroughly investigated in this study, particularly in the case of BTK inhibitors. Notably, the effects of IGFBP-3, Serpin E1 and PTX-3 are highly context- and cell-dependent^54–61^. IGFBP-3, for instance, has been shown to induce resistance to the PARP inhibitor olaparib in advanced prostate cancer^55^, but it sensitizes breast cancer cells towards antiestrogen resistance^56^. In line with our findings, PTX-3 has been associated with both cancer progression, therapeutic resistance ^57^, and inferior outcome in DLBCL patients^59^. However, it can also behave as a tumor suppressor^58^. Serpin E1, instead, has been implicated in promoting resistance in triple-negative breast cancer^60^ and non-small cell lung cancer^61^, and is associated with poor prognosis and cancer progression in various cancers^54^.

In line with our findings, several 3D BM stromal models have been developed to study the pathogenesis and therapeutic resistance in hematological malignancies^45^. For instance, in MM, 3D models mimicking interactions between MM cells and non-neoplastic BM cells have proven essential for understanding disease progression and drug resistance^62^. An interesting approach took advantage of the 3D Rotary Cell Culture System bioreactor, which allows myeloma-stroma co-cultures in gelatin scaffolds^63^. This model demonstrates that cell adhesion-mediated drug resistance was significantly higher in 3D cultures compared to 2D, and highlighted the role of the VLA-4/VCAM1 pathway in resistance to the proteasome inhibitors^63^. Similarly, the same model has been applied to study patient-specific responses to mobilizing agents in chronic lymphocytic leukemia^64^. Other advanced 3D models combine multiple BM niches to mimic the complex microenvironment supporting MM. A 3D bioprinting model incorporating both the endosteal and perivascular niches demonstrated the essential role of the perivascular niche in supporting MM cell proliferation^65^. In leukemias, 3D magnetic hydrogels seeded with mesenchymal stem cells (MSCs) can be used to create biomimetic BM environments, which result in greater resistance to chemotherapeutic agents compared to 2D cultures, further emphasizing the importance of 3D models for perhaps a more accurate drug testing^66^. A 3D decellularized bone scaffold model was used to investigate the role of ECM and MSC interactions in DLBCL. This model revealed that ECM and MSC interactions confer differential protection against doxorubicin-induced apoptosis depending on the lymphoma subtype, underlining the potential limitations of 2D cultures^67^.

Our 3D co-culture model distinguishes itself from existing literature due to its high reproducibility, scalability, and potential for drug screening applications. Moreover, our model has the potential to be integrated with further cellular components of the BM niche that might further sustain resistance, such as endothelial cells, osteoblast, and osteoclasts. Incorporating these cell types in future iterations could enhance the model, providing a more comprehensive and realistic representation of the BM lymphoma tumor microenvironment.

Despite the significant insights gained from this study, the specific survival pathways activated by the cytokines we identified are yet to be fully elucidated. Investigating these aspects will be critical for a deeper understanding of the resistance mechanisms. Furthermore, a direct comparison between our 3D model and *in vivo* models was not conducted, limiting our ability to evaluate how accurately it mirrors the drug responses observed in patients.

Even with these limitations, our 3D model allowed us to uncover potential novel mechanisms of microenvironment-driven resistance to BCR signaling inhibitors. Beyond its utility for dissecting the biological basis of drug resistance, our model holds significant clinical potential. With further optimization, its high-throughput nature could enable its use as a real-time drug screening platform, guiding therapeutic decisions for relapsed or refractory lymphoma patients.

## Acknowledgments

This work was supported by Fond’Action contre le cancer.

## Authorship

Contribution: A.Z., A.A., M.V.C., E.C. and A.A.M. performed the experiments; A.Z., A.A., L.C. analyzed the data and performed bioinformatic analysis; D.P. performed primary cell isolation; F.Si. provided mCherry positive cell lines; C.C. provided patient samples; F.Sp. and D.P. provided technical advice; A.Z., C.A., M.M. and F.B. designed the research and ideated the experiments; A.Z. wrote the manuscript; A.A., C.A., M.M. and F.B. revised the manuscript.

Conflict-of-interest disclosure: L.C. institutional research funds from Orion; travel grant from HTG; A.A. travel grant from Astra Zeneca, consultant for PentixaPharm; F.B. institutional research funds from ADC Therapeutics, Bayer AG, BeiGene, Floratek Pharma, Helsinn, HTG Molecular Diagnostics, Ideogen AG, Idorsia Pharmaceuticals Ltd., Immagene, ImmunoGen, Menarini Recherche, Nordic Nanovector ASA, Oncternal Therapeutics, Spexis AG; consultancy fee from BIMINI Biotech, Helsinn, Menarini; advisory board fees to institution from Novartis; expert statements provided to HTG Molecular Diagnostics; travel grants from Amgen, Astra Zeneca, Beigene, InnoCare, iOnctura. The other Authors have nothing to disclose.

