## Supplementary Appendix for "A High-Throughput Bone Marrow 3D Co-Culture System to Develop Resistance to B Cell Receptor Signaling Targeted Agents in B Cell Non-Hodgkin Lymphoma"

### Supplementary Materials and Methods

*Cell lines*

Established human cell lines derived from marginal zone lymphoma (VL51, K1718, SSK41, mCherry-VL51, mCherry-K1718 and mCherry-SSK41), diffuse large B-cell lymphoma (OCI-Ly-10 and TMD8), and mantle cell lymphoma (REC-1 and MINO) were cultured in RPMI 1640 medium supplemented with L-Glutamine (11875093, Gibco), 10% FBS and 1% P/S. mCherry-positive cell lines were established as previously described[27]. Cell line identity was authenticated by short tandem repeat DNA profiling, as previously described[28]. The cell lines were stored at -150°C, and all experiments were conducted within one to two months after thawing. Routine Mycoplasma testing was performed using the MycoStrip assay (rep-mysnc-100, InvivoGen) to confirm negativity.

*Supplementary Table 1*

Cell seeding concentrations and cytokine stimulation conditions used for MTT assays and Incucyte experiments as indicated in the *Material and Methods* section.

| Cell Line | Seeding Concentration (cells/well) | CHI3L1 | VCAM | DKK1 | IGFBP-3 | Serpin E1 | PTX-3 |
|---|---|---|---|---|---|---|---|
| | | Concentration (ng/mL), Time | | | | | |
| VL51 | $1 \times 10^4$ | 1, 4h | 1, 4h | 1, 4h | 1, 4h | 1, 4h | 1, 4h |
| Karpas1718 | $2 \times 10^4$ | | | | 10, 24h | 1, 24h | 100, 4h |
| SSK41 | $2 \times 10^4$ | | | | 10, 1h | 100, 24h | 100, 4h |
| OCI-Ly10 | $2 \times 10^4$ | | | | 1, 4h | 1, 1h | 1, 1h |
| TMD8 | $1 \times 10^4$ | | | | 1, 24h | 1, 4h | 1, 1h |
| REC-1 | $2 \times 10^4$ | | | | 100, 4h | 1, 1h | 100, 24h |
| MINO | $1 \times 10^4$ | | | | 100, 4h | 100, 4h | 100, 1h |

*Supplementary Table 2*

Drug concentrations used for MTT assays and Incucyte experiments as indicated in the *Material and Methods* section.

| Cell Line | Copanlisib | Ibrutinib |
|---|---|---|
| | Concentration (nM) | |
| VL51 | 10 | |
| Karpas1718 | | 1 |
| SSK41 | | 1 |
| OCI-Ly10 | | 0.5 |
| TMD8 | | 0.5 |
| REC-1 | | 2 |
| MINO | | 10 |

*Supplementary Table 3*

Antibody list used for IF as indicated in the *Material and Methods* section.

| Antibody | Catalog Number | Type | Host | Dilution Used |
|---|---|---|---|---|
| TLR4 Polyclonal Ab | bs-1201R | Primary | Rabbit | 1:200 |
| TGF beta R1 Polyclonal Ab | bs-0638R | Primary | Rabbit | 1:200 |
| Anti-TMEM219 | HPA059185 | Primary | Rabbit | 1:200 |
| uPAR recombinant mouse monoclonal Ab (2E2) | MA5-50263 | Primary | Mouse | 1:200 |
| Anti-Mouse Alexa Fluor™ 488 | A-28175 | Secondary | Goat | 1:1000 |
| Anti-Rabbit Alexa Fluor™ 488 | A-11008 | Secondary | Goat | 1:1000 |

*Supplementary Table 4*

Pannel of cytokines detected by cytokine array. Further details indicated in the *Material and Methods* section.

| Coordinate | Cytokine | Alias |
|---|---|---|
| A1-A2 | positive cnt | |
| A3-A4 | Adinopectin | |
| A5-A6 | Apolopoprotein 1 | |
| A7-A8 | Angiogenin | |
| A9-A10 | Angiopoietin-1 | |
| A11-A12 | Angiopoietin-2 | |
| A13-A14 | BAFF | |
| A15-A16 | BDNF | |
| A17-A18 | Complement component C5 | |
| A19-A20 | CD14 | |
| A21-A22 | CD30 | |
| A23-A24 | positive cnt | |
| B1-B2 | | |
| B3-B4 | CD40 ligand | |
| B5-B6 | Chitinase 3-like 1 | |
| B7-B8 | Complement Factor D | |
| B9-B10 | C-Reactive Protein | |
| B11-B12 | Cripto-1 | |
| B13-B14 | Cystatin C | |
| B15-B16 | Dkk-1 | |
| B17-B18 | CD26 | DPP4 |
| B19-B20 | EGF | |
| B21-B22 | CD147 | Emmprin |
| B23-B24 | | |

| | | |
|---|---|---|
| C1-C2 | | |
| C3-C4 | CXCL5 | |
| C5-C6 | ENG | CD105, endoglin |
| C7-C8 | TNSF6 | FAS ligand, CD178, CD95L |
| C9-C10 | FGF-2 | |
| C11-C12 | FGF-7 | |
| C13-C14 | FGF-19 | |
| C15-C16 | FLT3LG | FLT-3 ligand |
| C17-C18 | G-CSF | CSF3 |
| C19-C20 | MIC-1 | GDF-15 |
| C21-C22 | CSF2 | GM-CSF |
| C23-C24 | | |
| D1-D2 | CXCL1 | GROa, MSGA-a |
| D3-D4 | GH | Growth Hormone |
| D5-D6 | HGF | SF, Scatter Factor |
| D7-D8 | ICAM-1 | CD54 |
| D9-D10 | IFNG | IFN-gamma |
| D11-D12 | IGFBP-2 | |
| D13-D14 | IGFBP-3 | |
| D15-D16 | IL1a | IL-1a |
| D17-D18 | IL1b | IL-1b |
| D19-D20 | IL1ra | IL1-ra |
| D21-D22 | IL2 | IL-2 |
| D23-D24 | IL3 | IL-3 |
| E1-E2 | IL4 | IL-4 |
| E3-E4 | IL5 | IL-5 |
| E5-E6 | IL6 | IL-6 |
| E7-E8 | IL8 | IL-8 |
| E9-E10 | IL10 | IL-10 |
| E11-E12 | IL11 | IL-11 |
| E13-E14 | IL12 p70 | IL-12 p70 |
| E15-E16 | IL13 | IL-13 |
| E17-E18 | IL15 | IL-15 |
| E19-E20 | IL16 | IL-16 |
| E21-E22 | IL17a | IL-17A, CTLA8 |
| E23-E24 | IL18 Bpa | IL-18 Bpa |
| F1-F2 | IL19 | IL-19 |
| F3-F4 | IL22 | IL-22. IL-TIF |
| F5-F6 | IL23 | IL-23, IL-23A, SGRF |
| F7-F8 | IL24 | IL-24, C49A, FISP, MDA-7, MOB-5, ST16 |
| F9-F10 | IL27 | IL-27 |
| F11-F12 | IL31 | IL-31 |
| F13-F14 | IL32 | IL-32 |
| F15-F16 | IL33 | IL-33 |
| F17-F18 | IL34 | IL-34 |

| | | |
|---|---|---|
| F19-F20 | CXCL10 | IP-10 |
| F21-F22 | CXCL11 | I-TAC, SCYB9B |
| F23-F24 | KLK3 | Kallikrein, PSA |
| G1-G2 | Leptin | OB |
| G3-G4 | LIF | |
| G5-G6 | Lipocaliln-2 | NGAL, LCN2, Siderocalin |
| G7-G8 | CCL2 | MCP-1, MCAF |
| G9-G10 | CCL7 | MCP-3, MARC |
| G11-G12 | CSF1 | M-CSF |
| G13-G14 | MIF | |
| G15-G16 | CXCL9 | MIG |
| G17-G18 | CCL3 | CCL3/CCL4, MIP-1a/MIP-1b |
| G19-G20 | CCL20 | MIP-3a, Exodus-1, LARC |
| G21-G22 | CCL19 | MIP-3b, ELC |
| G23-G24 | MMP-9 | CLG4B, Gelatinase B |
| H1-H2 | MPO | Myeloperoxidase, Lactooperoxidase |
| H3-H4 | OPN | Osteopontin |
| H5-H6 | PDGF-AA | |
| H7-H8 | PDGF-AB/BB | |
| H9-H10 | PTX3 | Pentraxin 3, TSG-14 |
| H11-H12 | CXCL4 | PF4 |
| H13-H14 | RAGE | |
| H15-H16 | CCL5 | RANTES |
| H17-H18 | RBP-4 | |
| H19-H20 | RLN2 | Relaxin-2, RLXH2 |
| H21-H22 | Resistin | RETN, ADSF, FIZZ3 |
| H23-H24 | CXCL12 | SDF-1a, PBSF |
| I1-I2 | Serpin E1 | PAI-I, PAI-1, Nexin |
| I3-I4 | SHBG | ABP |
| I5-I6 | IL1R-L1 | IL-1 R4, IL1RL1, ST2, ST2L |
| I7-I8 | CCL17 | TARC |
| I9-I10 | TFF3 | ITF, TFI |
| I11-I12 | CD71 | TfR, TFR1, TFRC, TRFR |
| I13-I14 | TGF-a | TGFA |
| I15-I16 | Thrombospondin-1 | THBS1, TSP-1 |
| I17-I18 | TNF-a | TNFSF1A |
| I19-I20 | uPAR | PLAUR |
| I21-I22 | VEGF | BEGFA |
| I23-I24 | | |
| J1-J2 | positive cnt | |
| J3-J4 | | |
| J5-J6 | Vitamine D BP | VDB, DBP, VDBP |
| J7-J8 | CD31 | PECAM-1 |
| J9-J10 | TIM-3 | HAVCR2 |
| J11-J12 | VCAM-1 | CD106 |

| | |
|---|---|
| J13-J14 | |
| J15-J16 | |
| J17-J18 | |
| J19-J20 | |
| J21-J22 | |
| J23-J24 | negative cnt |

*Supplementary Table 5*

Ibrutinib $IC_{50}$ calculated in 2D for the different cell lines.

| **Cell line** | **Ibrutinib IC50 (nM)** |
|---|---|
| Karpas1718 | 122 |
| SSK41 | 788 |
| VL51 | 950 |
| REC-1 | 15 |
| MINO | 132 |
| OCI-Ly10 | <10 |
| TMD8 | <10 |

### Supplementary Figures

#### Supplementary Figure 1

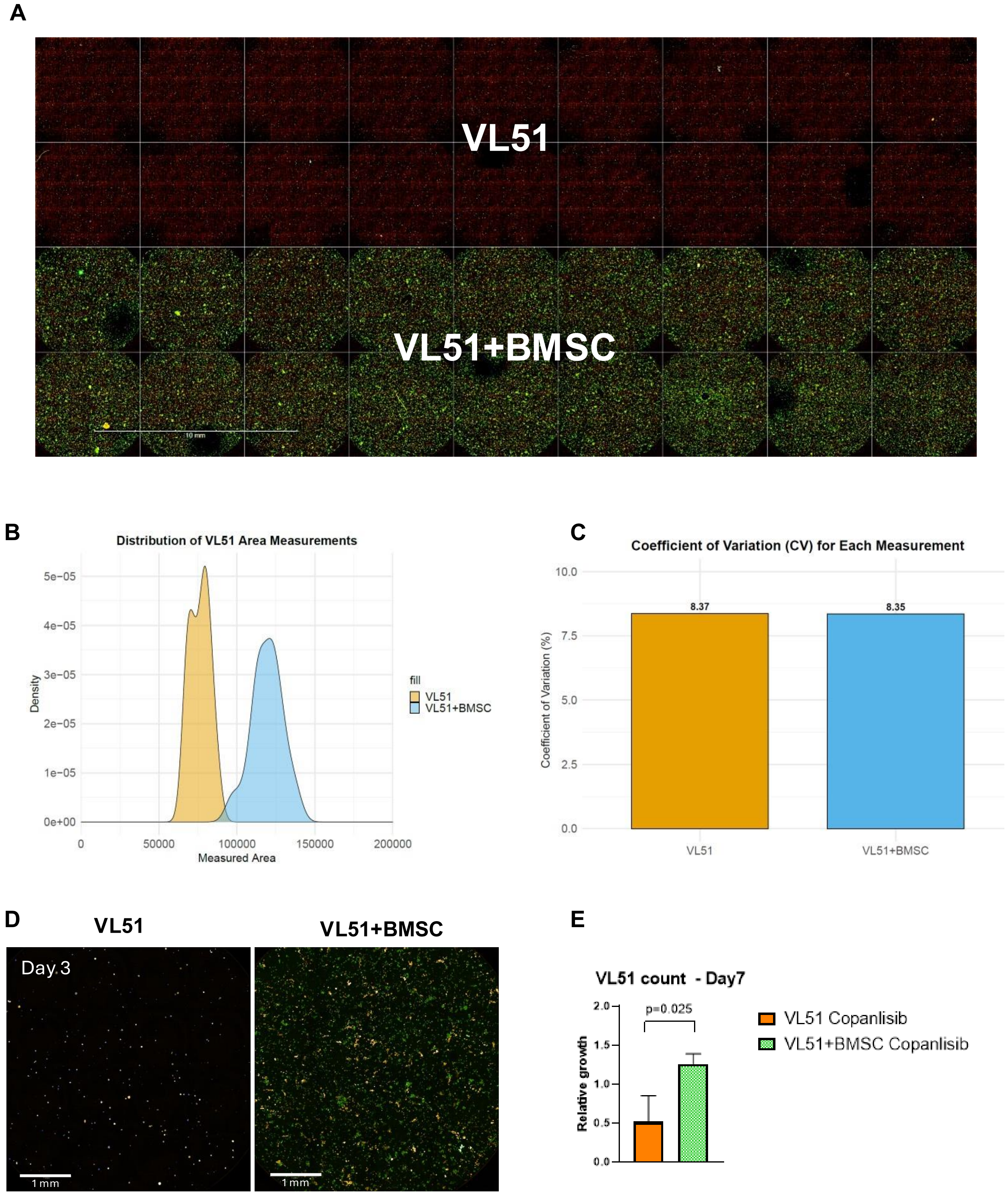


#### Supplementary Figure 1

**(A)** Representative immunofluorescence maximum projection images of VL51 (in orange) cultured in presence or absence of BMSCs (in green) on day 0 post seeding. Images taken with 10x Air objective NA 0.3, WD 5.2mm of the Opera

Phenix Plus High-Content Screening System while keeping the plate at 37°C, 5% $CO_2$, and optimal humidity.

**(B)** Plot showing the distribution of the VL51 measured areas on maximum projection images (Supplementary Figure 1A) between wells and among mono- (orange) and co-culture (light blue) conditions. Plot generated with R.

**(C)** Bar plot showing the coefficient of variation of the VL51 measured areas on maximum projection images (Supplementary Figure 1A) between wells and among mono- (orange) and co-culture (light blue) conditions. Plot generated with R.

**(D)** Representative immunofluorescence maximum projection images of VL51 (in orange) culture in the presence or absence of BMSCs (in green) on day 3 before treatment. Images taken with 10x Air objective NA 0.3, WD 5.2mm of the Opera Phenix Plus High-Content Screening System while keeping the plate at 37°C, 5% $CO_2$, and optimal humidity.

**(E)** Bar plot showing VL51 count on maximum projection images of VL51 (in orange) culture in presence or absence of BMSCs (in green) upon DMSO or 100nM copanlisib treatment on day 7. Images taken with 10x Air objective NA 0.3, WD 5.2mm of the Opera Phenix Plus High-Content Screening System while keeping the plate at 37°C, 5% $CO_2$, and optimal humidity. Statistical significance tested with t test.

**Supplementary Figure 2**

**A**

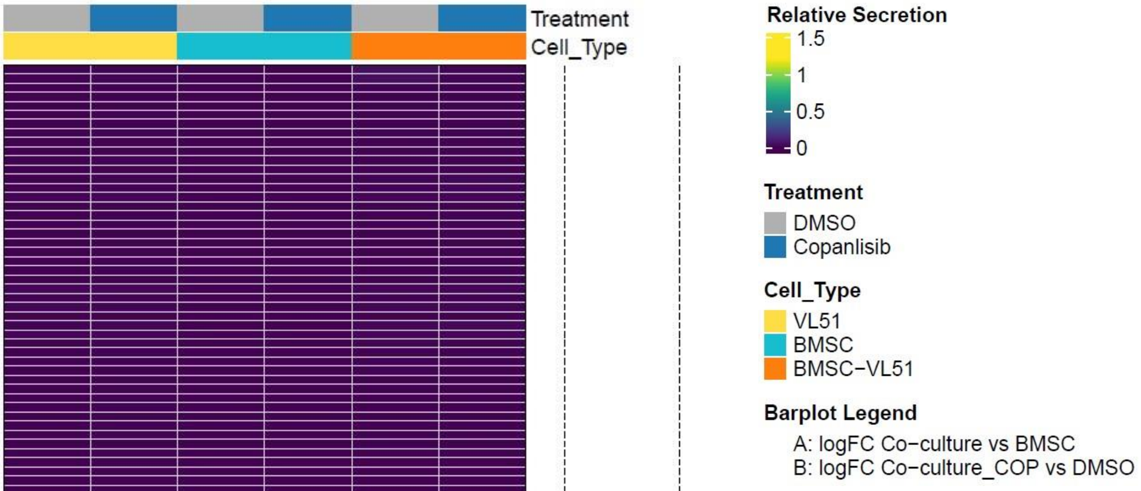


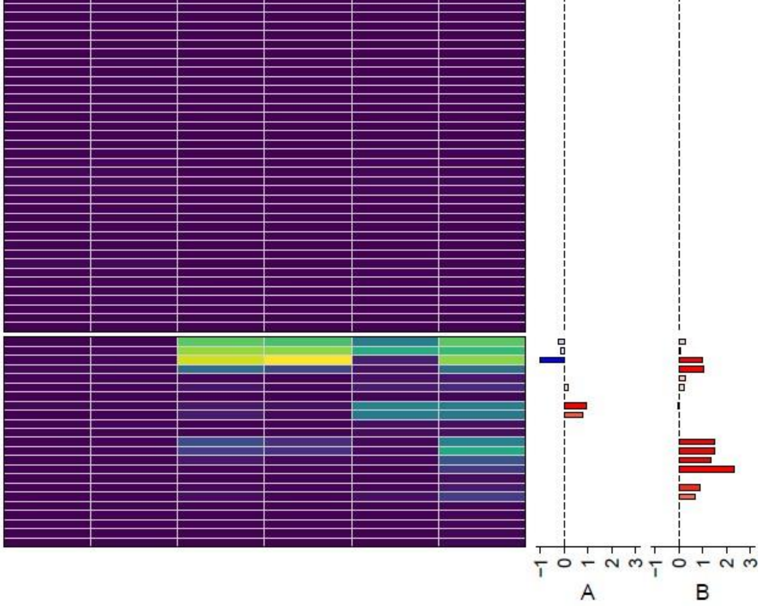


**C**

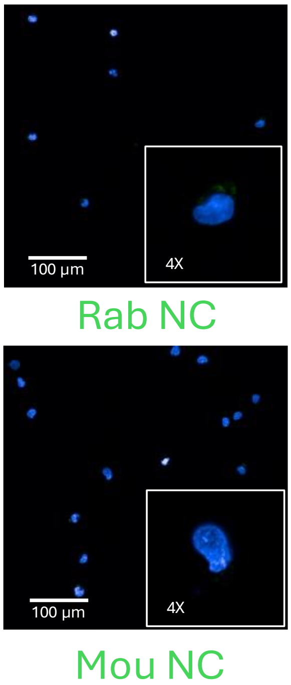


**B**

| | VL51 | BMSC | VL51+BMSC |
|---|---|---|---|
| DMSO | 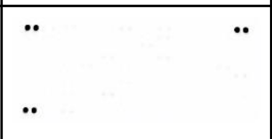 | 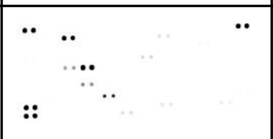 | 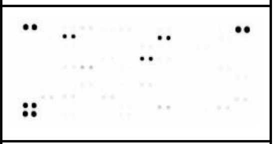 |
| Copanlisib | 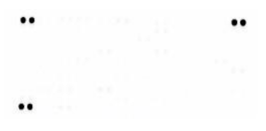 | 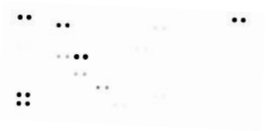 | 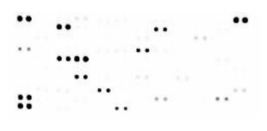 |

**Supplementary Figure 2**

- **(A)** Plot showing the whole panel of cytokines analyzed with the cytokine array. The heatmap was generated with the relative secretion values (row values normalized to internal positive and negative controls) of each cytokine in the different culture conditions. The bar plots show the log fold change values comparing co-culture to mono-culture upon DMSO (A) or the copanlisib to the DMSO conditions in the co-culture (B).
- **(B)** Representative images of the cytokine array membranes of all conditions.
- **(C)** Secondary antibody control for VL51 cytokine receptor IF images. In green it is shown the expression of the specific secondary antibody signal, in blue the nuclei. Images taken with 40x Water objective NA 1.1, WD 0.62mm of the Opera Phenix Plus High-Content Screening System at room temperature. (Rab=anti-rabbit secondary antibody; Mou=anti-mouse secondary antibody)

**Supplementary Figure 3**

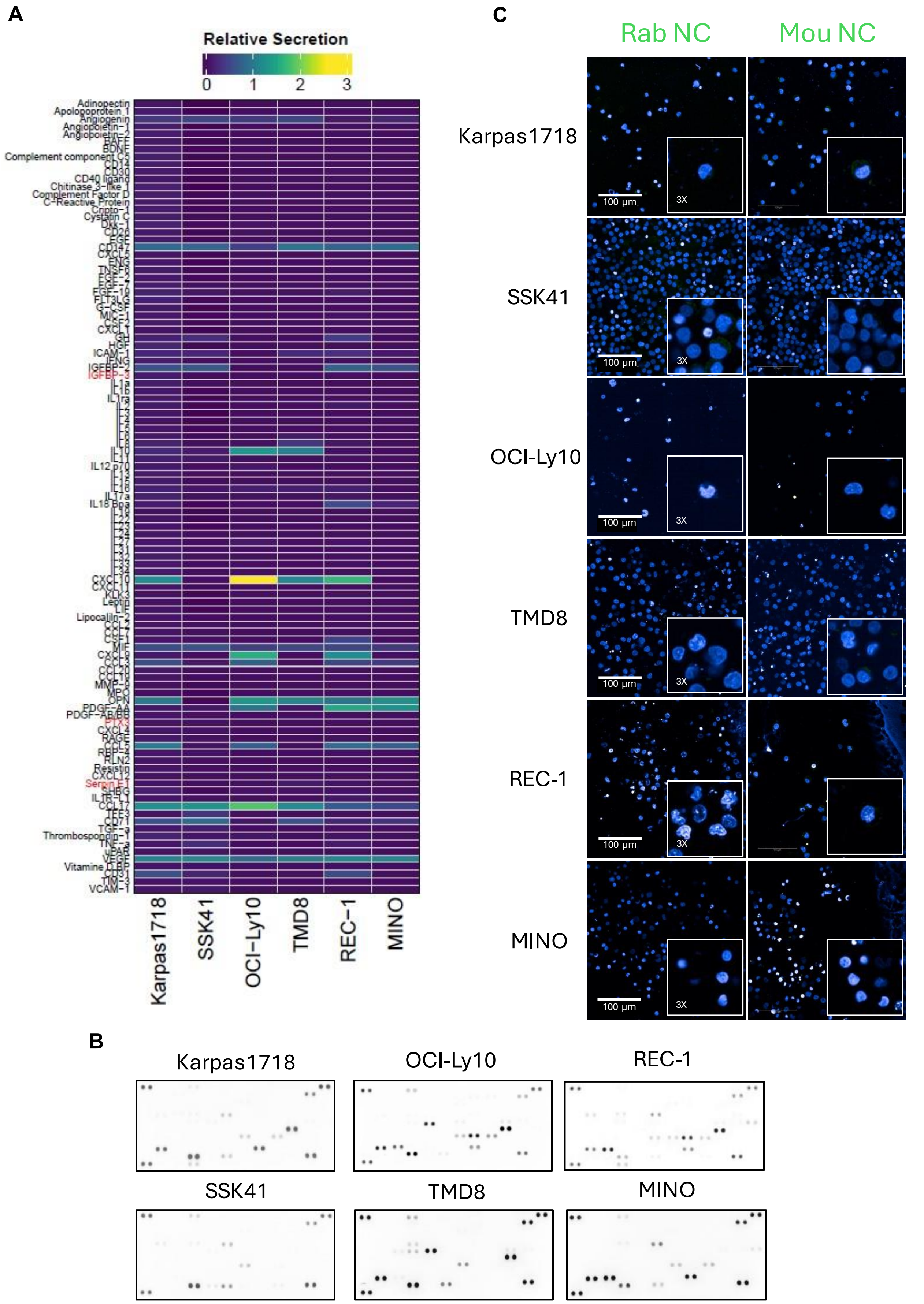

**Supplementary Figure 3**

**(A)** Plot showing the whole panel of cytokines analyzed with the cytokine array. The heatmap was generated with the relative secretion values (row values normalized to internal positive and negative controls) of each cytokine in the different culture conditions. The bar plots show the log fold change values comparing co-culture to mono-culture upon DMSO (A) or the copanlisib to the DMSO conditions in the co-culture (B).

**(B)** Representative images of the cytokine array membranes of all conditions.

**(C)** Secondary antibody control for the indicated cell line cytokine receptor IF images. In green, it is the expression of the specific secondary antibody signal, in blue, the nuclei. Images taken with 40x Water objective NA 1.1, WD 0.62mm of the Opera Phenix Plus High-Content Screening System at room temperature. (Rab=anti-rabbit secondary antibody; Mou=anti-mouse secondary antibody)

**Supplementary Figure 4**

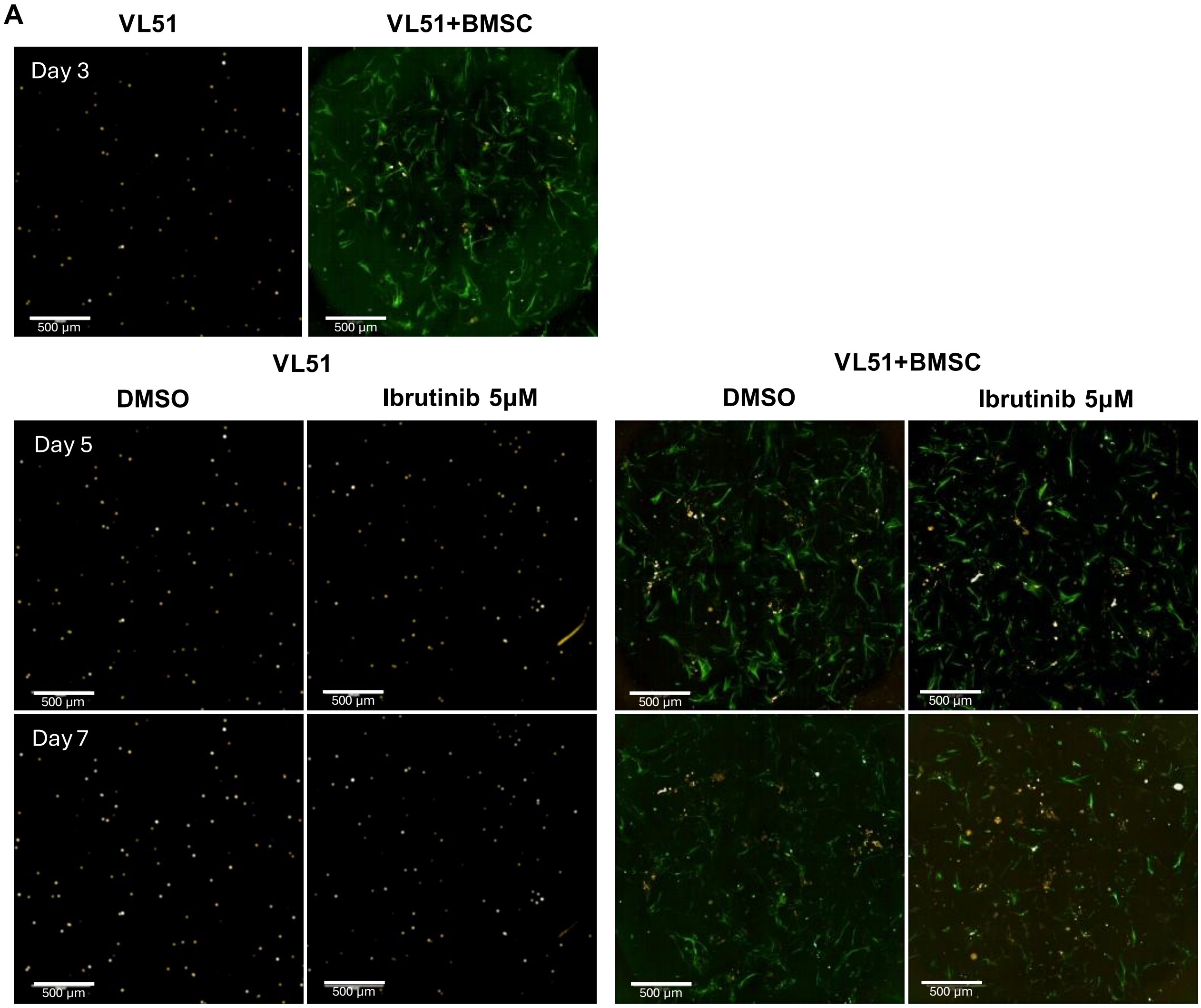

**Supplementary Figure 4**

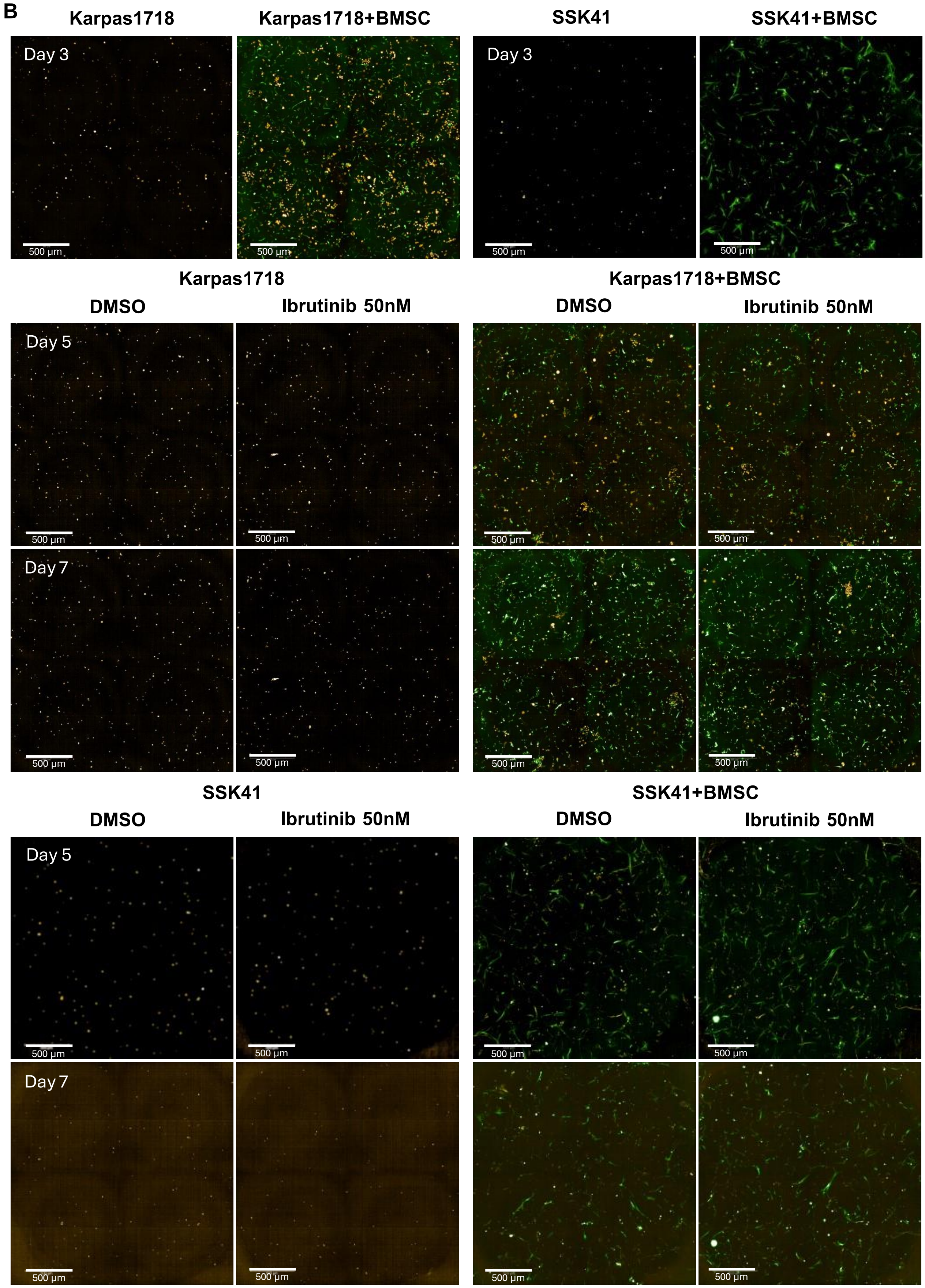

**Supplementary Figure 4**

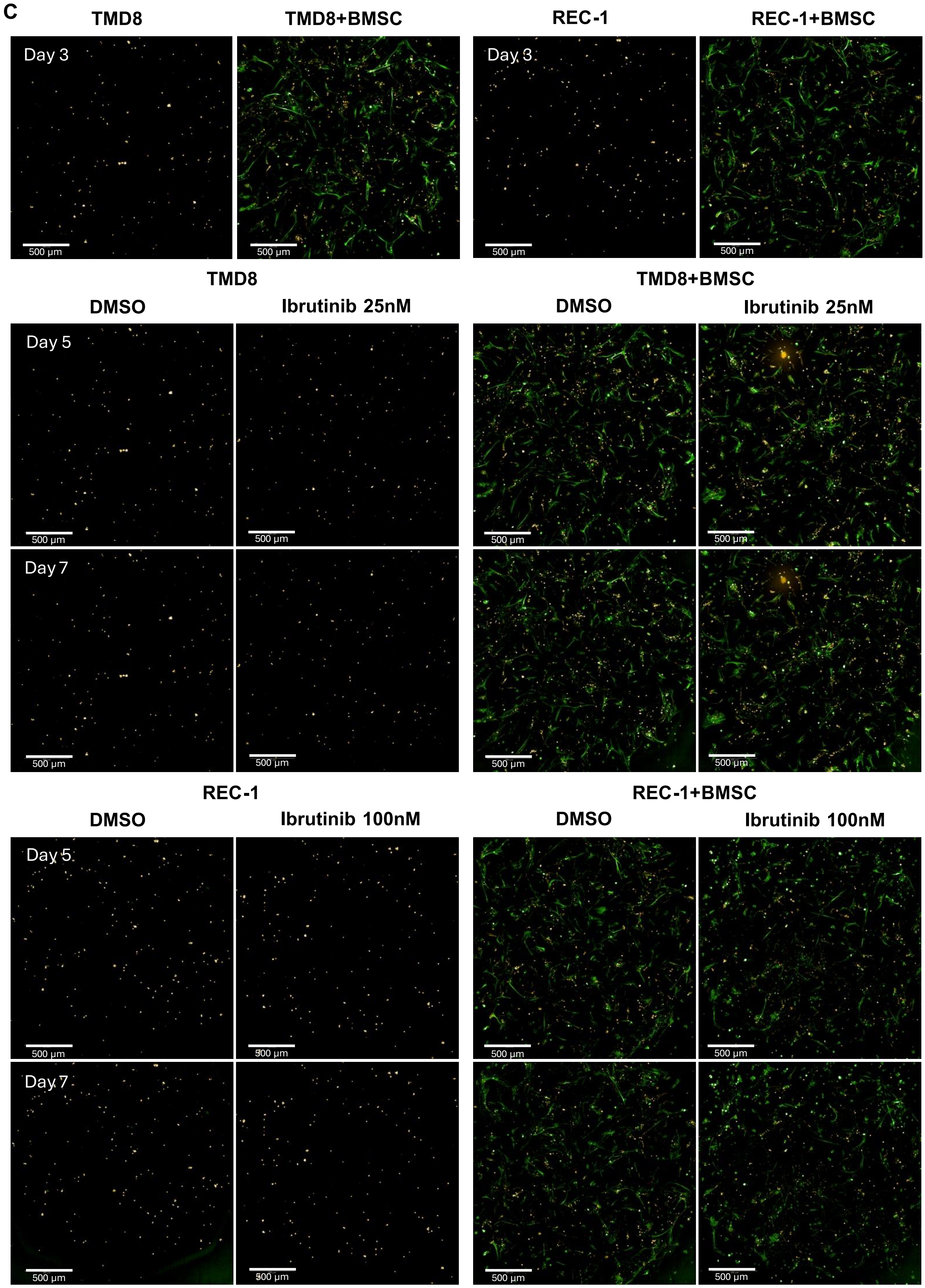

**Supplementary Figure 4**

**(A-C)** Representative immunofluorescence maximum projection images of the indicated B cell lymphoma cell lines (in orange) cultured in the presence or absence of BMSCs (in green) on day 3, 5, and 7 upon indicated culture conditions. Images taken with 10x Air objective NA 0.3, WD 5.2mm of the Opera Phenix Plus High-Content Screening System while keeping the plate at 37°C, 5% $CO_2$, and optimal humidity.
